# Targeted chelation therapy decreases NLRP3 expression by vascular cells and acts as senomorphic in Chronic Kidney Disorder induced Vascular Calcification

**DOI:** 10.1101/2025.04.23.650230

**Authors:** Shivani Arora, Gregory Halsey, Fatema-Tuj Zohora, Alyssa Swiss, Narendra Vyavahare

## Abstract

**Background:** Chronically high levels of phosphate (P) in the serum caused by chronic kidney disease (CKD) induce osteogenic changes in the aortic Vascular Smooth Muscle Cells (VSMCs). Premature onset of cellular senescence is observed in these phenotypically transitioned cells, which plays a critical role in pathology of vascular calcification. We have previously shown that EDTA therapy can remove calcium deposits from the arteries in a rat model of CKD and reduces the expression of osteogenic markers in the aorta. In the current study we evaluated if chelation therapy with EDTA has senotherapeutic potential and could also decrease the accumulation of senescent cells in the aorta once it has calcified.

**Methods:** We used an adenine diet-based rodent model of late-stage CKD and an ex-vivo aortic ring culture model to evaluate the senotheraputic potential of EDTA loaded-human serum albumin nanoparticles tagged with anti-elastin antibody-Flexibzumab (EDTA-NP). For validation we performed a comparative proteomics analysis on the total proteins harvested from the abdominal aortas of the EDTA nanoparticle treated and untreated animals.

**Results:** Our results show that targeted chelation therapy with EDTA-NP decreases the percentage of SA-beta gal positive senescent cells in the calcified aorta and acts as senomorphic by decreasing NLRP3 inflammasome formation which is a major intracellular source of Senescence associated secretory phenotype (SASP).

**Conclusion:** For the first time, the current study provides a proof of concept on the senotheraputic potential of a targeted chelation therapy and its capacity to modulate SASP from the senescent cells accumulates in calcified aorta.

**Highlights:**

- Our findings show that chelation therapy can act as senomorphic, and increases the life span of rodents suffering from heavy vascular calcification.
- Chelation therapy decreases senescent cell accumulation, SASP and NLRP3 expression in the aorta.
- Chelation therapy is a novel method for reprogramming senescent cells in the aorta to prevent their phenotypic switching to inflammatory senescent cells and ultimately to osteoblasts.
- Current data have provided a new hypothesis that agents that restore mineral imbalance in the cellular microenvironment (in this case, EDTA) have the potential to act as senomorphics, which can serve as safer therapeutic alternatives over senolytics to treat vascular calcification by decreasing apoptosis.

## Introduction

Aging-associated medial artery calcification (MAC), also known as Mönckeberg’s calcification, is the most common form of calcification that is observed during aging, and it is a major contributor to cardiovascular morbidity and mortality. The aging process is typically accelerated in patients with chronic kidney disorder (CKD), as is evidenced by progressive vascular disease, persistent uremic inflammation, muscle wasting, osteoporosis, and frailty that precedes terminal renal failure (1,2). Although the exact underlying mechanisms of accelerated vascular aging in CKD have not yet been fully elucidated, many preclinical studies emphasize on the critical role played by the increased number of senescent cells that accumulate in the calcification of aorta. Recently, Fang et al reviewed the functional role of cellular senescence during vascular calcification in CKD (3). Oh et al (4) have demonstrated the presence of senescent cells (as defined by p16^High^ and p53 positive expression) in the aortic valves of individuals suffering calcified aortic valve disease. These results indicate that the severity of tissue remodeling in CKD and in calcific aortic valve disease is directly proportional to the percentage P16^INK4A^ expressing cells. Preclinical data from Judith Campisi and James Kirkland have supported the view that senolytics -agents that cause selective apoptosis of senescent cells-can be potential therapy for treating vascular stenosis by clearing senescent cells from atherosclerotic plaques (5–7). These studies support the hypothesis that senolytics and senomorphics (agents that alter SASP and modulate the local immune environment) could provide an attractive therapeutic strategy to prevent and treat cardiovascular diseases.

Vascular calcification is accompanied by severe pro-inflammatory response that exacerbates the progression of disease (8,9). The highly inflammatory secretory phenotype from senescent (SASP) cardiovascular cells has been reported to induce osteogenic changes in the aortic VSMCs (10–12). NLRP3 inflammasome activation in the cardiac cells has been identified as key source of cellular senescence and SASP leading to premature cardiac aging and progression of cardiovascular disorders (13–15).

Multiple upstream pathways have been reported to cause activation of NLRP3 inflammasome signaling during vascular calcification, direct stimulation by Calcium crystals is one of the most direct and important mechanism that causes activation of NLRP3 inflammasome in the calcified aorta. Upon activation NLRP3 inflammasome undergoes self-oligomerization and rescues apoptosis-associated spec-like protein (ASC) from degradation. ASC is a death domain family protein with a Caspase 1 Activation and Recruitment domain. ASC rescue turns on the caspase-1 effector pathway, leading to conversion of pro IL-1β to IL-1β, and pro-caspase-1 to caspase-1 and ultimately the release of active caspase 1, IL-1β and IL-18 causing amplification of inflammatory responses as well as pyro-apoptosis (16–19).

We have previously demonstrated that chelation therapy with EDTA loaded nanoparticles that specifically target degraded elastin reversed existing heavy mineral deposits in arteries. The reversal of aortic calcification was accompanied by a significant reduction of bone-associated mRNA expression of *BMP2* and *RUNX2,* reflecting that removal of calcium deposits by targeted chelation has the potential to prevent phenotypic transition of VSMCs to osteoblasts at genetic level (20,21). BMP2 is a key transcription factor that regulates the expression of both p21 and RUNX2 at transcriptional level (22). Prompted by this preclinical evidence, for the current study we analyzed the impact of EDTA-based chelation therapy on the senescence markers and the NLRP3 expression, identified as a potential source of SASP(23–25).

We employed an in-vitro aortic ring culture model and in-vivo chronic kidney disease model to investigate the impact of targeted chelation therapy on the status of senescent cells in the calcified aorta. The samples were tested for the presence of senesce, and SASP markers, as well as for NLRP3 expression in the aorta. Human serum albumin nanoparticles loaded with EDTA were conjugated with Flexibzumab antibody that can specifically target damaged elastin in the artery (21,26,27) were injected *iv* in the late-stage CKD animals, and aortas from the animals were harvested to analyze calcification status, senescence phenotype, and NLRP3 expression. Overall, this study provides a proof of concept that targeted chelation therapy not only reverses calcification but also can act as senotheraputics to mitigate cellular senescence observed in vascular calcification.

## Material And Methods

### Materials

Male Sprague Dawley rats (Charles River, CD® IGS strain code-001), Adenine diet containing 2.5% protein (Envigo Teklad Custom Diet TD. 130127, Madison, WI), EDTA (Sigma, E6511), vitamin D3 (VitD3, Cholecalciferol, Sigma, #C9756-5G), NaCl (sigma, S3014), KCl (Sigma P9541), NaHCO3 (Sigma, S5761), glucose (Sigma, G7021), NaH2PO4 (Sigma, S0876), dPBS (Gibco, 10010-031), DMEM (Gibco-1885-076), FBS, Pen-Sterp, Human Aortic Smooth Muscle Cells (SMCs) (Promocell, C12533), SMC Medium II (Promocell 22062), ABT263, HSA (Evonik Birmingham Laboratory, 777HSA097), HEPES (Sigma, H3375), absolute ethanol (Fisher BP 2818500), mPEGNHS, MW 2000 (BOC Sciences), Traut’s reagent (G-Biosciences, Saint Louis, MO, BC95), 5-Dodecanoylaminofluorescein di-β-D-Galactopyranoside (C12FDG, MedChem Express HY-126839), Cell Staining buffer (BioLegend 00-4222-26), anti-CD16/32 mouse monoclonal antibody (BioLegend Cat#156604), anti-CD45-APC (eBiosciences Cat#17-0461-82) DAPI (BioLegend Cat#422801), Rabbit anti-Rat Caspase-3 (Cell Signaling Technology, C8487), NLRP3 (Novus Biologicals, NBP2-12446), and PiT-1 (Santa Cruz Biotechnology,SC98824), anti-rabbit IgG secondary antibody Cy5 (Invitrogen, A10523) cDNA synthesis Kit (Bio-Rad, Cat# 1725034), iTaq-Universal SYBR Green Super mix 12 reagent (Bio-Rad, Cat# 172512), SA-βGal Staining Kit (Cell Signaling, 9860), Rat IL-6 ELISA Kit (RnD SYSTEMS DY-506), Rat IL-1β ELISA Kit (RnD SYSTEMS SRLB00), T-Per protein extraction buffer (Thermo Fischer, 78510), BCA assay kit (Thermo Fisher Scientific,A55865).

### Animals

Male Sprague Dawley rats age (12 weeks old) were used for the study. This study was approved by the Institutional Animal Care and Use Committee (IACUC) at Clemson University (Animal Use Protocol number 2021-006). The study was carried out in compliance with the ARRIVE guidelines. Animals obtained from Charles River Laboratories were acclimatized for two weeks before starting the study and were maintained on a standard rodent diet (Teklad Global 18% Protein Rodent Diet, Madison, WI). During the entire study, the animals were monitored for body weight, temperature, and regular activities by an experienced veterinarian and were euthanized by saline perfusion under isoflurane anesthesia when they reached a humane endpoint of >20% weight loss.

### Early-Stage vascular calcification in Chronic Kidney Disorder (CKD) Model

The animals were randomly divided into treatment and control groups. Control group animals were maintained on a standard chow diet till the completion of the study, and the animals in the treatment groups were maintained on 0.75% Adenine diet containing 2.5% protein with a modified schedule for 28 days as described previously (28) The animals were euthanized by saline perfusion under isoflurane anesthesia, and the aortas (from the heart to the iliac bifurcation), as well as other organs, including lungs, liver, kidneys, and spleen, were harvested and preserved accordingly for further analysis. Blood was also collected by cardiac puncture for serum analysis (Supplementary Figure 1A).

### Late-Stage heavy vascular calcification in CKD Model

The animals were randomly divided into treatment and control groups. Control group animals were maintained on a standard chow diet till the completion of the study, and the animals in the treatment groups were maintained on a 0.75% Adenine diet containing 2.5% protein with a modified schedule for 28 days. Five days after completion of the adenine diet, the animals in the treatment groups were either injected intraperitoneally with vitamin D3 (8.75 mg/kg/day formulated in olive oil) or equivalent olive oil as a vehicle for four consecutive days. 3 days post I.P injections the animals in the treatment group that received vitamin D injections were further randomly subdivided into two groups: (i) the EDTA -NP group – in which the animals received two weekly injections (a total of 5 injections) of EDTA loaded nanoparticles conjugated with elastin targeting antibody (10mg/kg IV), and (ii) the control group – in which the animals received blank nanoparticles conjugated with elastin targeting antibody (Blank-NPs) with the same dosage as EDTA nanoparticles. After three weeks of nanoparticle therapy, the aortas (from the heart to the iliac bifurcation) and other organs, including lungs, liver, kidneys, and spleen, were harvested and preserved accordingly for further analysis. Blood was also collected for serum isolation and biomarker analysis (Supplementary Figure 1B).

### Aortic Ring Culture for vascular calcification

Male Sprague-Dawley rats (16 weeks old) were euthanized by saline perfusion under isoflurane anesthesia and full-length aortas were harvested. under the aseptic condition in ice-cold Moscona’s buffer (8 g NaCl, 0.2 g KCl, 1 g NaHCO3, 1.7 g glucose, 0.005 g NaH2PO4 in 1 L DI water; pH 7.2). After washing four times with Dulbecco’s phosphate buffered saline (PBS), the aortas from ascending to the iliac bifurcation were cut into pieces to obtain 1cm aortic rings. These rings were then cultured in Dulbecco’s Modified Eagle Medium (DMEM) with 1% heat-inactivated FBS and 1% penicillin-streptomycin in a humidified incubator at 37◦C, under 5%CO_2_ for overnight equilibration. The aortic rings were treated with 0.5U/ml elastase for 40 minutes at 37^◦^C to induce initial elastin damage. Subsequently, elastase-treated aortic rings were washed twice with PBS and incubated in high phosphate medium in DMEM (final concentration of Pi: 2.9mM) containing 1% heat-inactivated FBS and 1% penicillin-streptomycin for ten days. The rings were then cultured with 0.5mg/ml pure EDTA (corresponding to the same concentration as delivered in vivo with HSA nanoparticles) in the presence of high phosphate in DMEM and 10% FBS for three days. The experimental groups were: (i) DMEM only (0.9mM phosphate; denoted as a control group); (ii) elastase & high phosphate treated with no EDTA (high Pi group); (iii) elastase & high phosphate treated followed by EDTA (EDTA-treated group). NaH_2_PO_4_.2H_2_O in a final concentration of 2.9 mM was used as source of Pi. During incubation, the conditioned medium was replaced every two days (Supplementary Figure 2).

### Vascular smooth muscle cell calcification studies

Human aortic smooth muscle cells (HAoSMCs) were purchased from Promocell and used between passages four and five for all experiments. Monocultures were maintained in 75 cm^2^ tissue culture-treated vented flasks in a 37°C and 5% CO_2_ environment in smooth muscle cell Medium II (Promocell). To create high phosphate conditions, NaH_2_PO_4_.2H_2_O was added to obtain a final concentration of 2.9 mM of Pi. EDTA was added to achieve the final concentration of 0.5 mg/ml in the EDTA treatment group in the culture media. ABT263 (1 mM) was used as known senolytic agent for comparison.

### EDTA Np Preparation and Conjugation with Flexibzumab Antibody

The desolvation method was used to make EDTA-loaded human serum albumin (HSA) nanoparticles following the method described in recently published manuscript from our group (20). Briefly, 200 mg of HSA (SeraCare, Milford, MA) was dissolved in 4ml of deionized water, and 50 mg of disodium salt of EDTA (Fisher Scientific, NJ) was then dissolved in HSA solution. The pH of the solution was adjusted to 8.5. The solution was then added dropwise to the absolute ethanol (1 mL/min) under constant stirring, followed by the addition of 25 µl of 8% glutaraldehyde as a crosslinker. The solution was incubated at room temperature for two hours with constant stirring at 800 rpm. Nanoparticles thus formed were centrifuged at 6000rpm for 10 minutes, rinsed in deionized water (saturated with EDTA), and resuspended in phosphate-buffered saline before conjugating with thiolated anti-elastin antibody conjugation (29). 10 mg of formulated nanoparticles were PEGylated with 2.5 mg of α-maleimide-ω-N-hydroxysuccinimide ester poly (ethylene glycol) for an hour at room temperature under gentle agitation. Meanwhile, 20 µg of custom-made humanized anti-elastin antibody was added to 68 μg of Traut’s reagent for antibody thiolation, and subsequently the mixture was incubated in 4-(2-hydroxyethyl)-1-piperazineethanesulfonic acid (HEPES) buffer (20 mM, pH=8.8) at room temperature for an hour under gentle agitation. Finally, thiolated antibody was added to the PEGylated nanoparticles and incubated overnight (16 hours) at 4^0^C under gentle rocking for conjugation.

### SA-Beta Galactosidase Activity

Cells isolated from the abdominal aorta were counted and adjusted to 3×10^6^ cells/ml in complete culture media, sterile DMEM containing DNase, and incubated with 66 mM C12FDG for 1 hour in a 37°C water bath. Cells were then spun down at 500 g for 5 minutes, washed with Cell Staining buffer (BioLegend), blocked with anti-CD16/32 mouse monoclonal antibody (BioLegend Cat#156604), and stained with anti-CD45-APC (eBiosciences Cat#17-0461-82) each at a 1:100 and 1:200 dilution on ice for 30 minutes. Cells were then washed in Cell Staining buffer and resuspended in Cell Staining buffer containing 1X DAPI (BioLegend Cat#422801). Samples were analyzed on an Attune acoustic focusing cytometer. Gates were set as follows: cells (FSC-A/SSC-A), forward scatter singlets (FSC-H/FSC-A), side-scatter singlets (SSC-H/SSC-A), live cells (DAPI-ve), non-immune cells (CD45-ve), and were analyzed for C12FDG fluorescence. Data were analyzed with FlowJo (v.10)(30).

### Immunohistochemistry for Caspase 3, NLRP3, and PiT-1

Abdominal aortas with cryosections of 7 µm thickness were fixed in cold acetone for 10 minutes, followed by rehydration along with decalcification in 0.5% of disodium ethylene diamine tetra acetic acid in Dulbecco’s phosphate buffered saline (PBS) for 15 minutes at room temperature. The sections were then washed in PBS and subsequently blocked with Background Buster for 30 minutes at room temperature and incubated at 4^◦^C overnight with primary antibodies: Rabbit anti-Rat Caspase-3 (Cell Signaling Technology, Danvers, MA), NLRP3, and PiT-1 (Santa Cruz Biotechnology, Dallas, Texas). Stained sections were then incubated with appropriate secondary antibodies conjugated with either anti rabbit Cy3 or Cy5 IgG for visualizing the target proteins. Finally, the aortic sections were counterstained with DAPI and mounted with an aqueous mounting medium for imaging. Four to six sections from equal spacings were taken from each sample for the semiquantitative analysis of target proteins.

### Reverse transcriptase quantitative PCR for gene expression

The expression of genes of interest was performed in aorta samples. The tissue was stored immediately after retrieval in the Trizol. The tissue was homogenized in Trizol using Powergen 125 (FS-PG125, Fisher Scientific) homogenizer, and RNA was extracted using the phenol-chloroform extraction method. A nanodrop instrument was used to analyze the extracted RNA through quantitative and qualitative analysis. cDNA synthesis was performed using 200ng of total RNA with iScript gDNA clear cDNA synthesis Kit (Bio-Rad, Cat# 1725034). The total cDNA obtained was diluted five times before amplification with iTaq-Universal SYBR Green Supermix 12 reagent (Bio-Rad, Cat# 172512) on Bio-Rad quantitative PCR platform (96-well format). Quantification was performed using the delta-delta CT method, using GAPDH as a housekeeping gene. A list of Primers used (OCN, RUNX2, IL-6, IL-1β, MCP-1, p21, p19, and GAPDH) in the study is given in supplementary Table-1.

### IL-6 and IL-1β measurements by ELISA

Culture supernatants were centrifuged at 2000 × g for 10 min at 4 °C to remove cellular debris. Cytokine concentrations in cell culture supernatants and animal serum were analyzed using ELISA kits (Invitrogen) according to the manufacturer’s protocol.

### MMP activity analysis using Zymography

MMPs activity in the serum was measured using in-gel zymography, with slight modifications(31). Briefly, serum samples from the treatment group (containing 20 µg of total protein) were loaded on gelatin-impregnated SDS gels (ZY00105, Fisher Scientific). Coomassie blue staining was used to visualize the sites of proteolytic digestion.

### Microcomputed tomography (microCT) imaging for calcification

Calcification or mineral deposits in rat aortas and kidneys were scanned using microCT (Bruker Skyscan 1176, Billerica, MA). Immediately after sacrifice, rat aortas were explanted and kept in cold PBS in 50 ml tubes for micro CT scanning. Aortic scanning was performed using a 0.5mm Aluminum filter at a voltage of 90kV and a current of 278 µA, with 45ms exposure time. Aorta specimens were scanned using a 360^◦^ rotation with a step of 0.7^◦^. The reconstruction of x-ray back-projection images into cross sections was performed using Bruker’s NRecon Software, which uses modified Feldkamp’s algorithms. The range of attenuation coefficients was the same across all aortas to 10, acquiring a comparable result between blank Nps and EDTA-NPs groups. For comparative analysis, Bruker’s volume rendering CTVox software was used to represent reconstructed images as a 3D object, with the same settings over aortas or kidneys. Further, 3D morphometric analysis was conducted with CTAn software from Bruker.

### Alizarin Red Staining for calcification

Optimal cutting temperature (OCT) compound– embedded frozen sections (7µm each) were stained with 2% Alizarin Red solution (pH 4.1-4.3) for 1 minute. The stained sections were then washed with two changes of distilled water for 10 minutes each and subsequently dehydrated in graded alcohol and xylene for mounting and bright field imaging with Keyence BZ-810. Semiquantitative analysis for percentage positive stain was performed with equally distributed four to six serial sections from each abdominal aortic sample.

### Statistical Analysis

Statistical analysis was performed using GraphPad Prism 10. Results were analyzed by One way ANOVA test. The number of biological replicates (n) is indicated in the figure legend. Statistical comparisons were performed between groups of rats, and/or samples, using two-tailed t tests for two groups, or one-way ANOVA, for three, with corrections for multiple testing. p < 0.05 was considered significant.

### Proteomics Analysis

A 3 cm section of abdominal aorta from CKD-VitD model rats was homogenized, and protein was isolated using T-Perm protein extraction buffer as per manufacturer’ protocol (Thermo Fischer). Protein concentrations were determined using BCA assay kit (Thermo Fisher Scientific, Watham, MA, USA). Protein samples were normalized to 60 μg with MS-grade water and proteins were reduced with 20 mM tris(2-carboxyethyl) phosphine (TCEP) by incubating at 50°C for 15 minutes. Proteins were brought to room temperature and then alkylated with 40 mM iodoacetamide (IAA) by incubating in dark at room temperature for 30 min. Tryptic digestion was performed using suspension traps (S-trap mini, Protifi, Fairport, NY, USA) following the manufacturer’s protocol. The reduced and alkylated proteins (60 μg) were acidified with 10:1 sample/12% phosphoric acid (v/v) and then diluted with 1:7 acidified sample/Binding Buffer (v/v). Proteins were loaded to S-traps in aliquots of 200 μL, centrifuged at 4,000 g for 30 sec, discarding the flow-through, washed six times with 200 μL Binding Buffer, discarding the flow-through, and centrifuged at 4,000 g for 1 min. Trypsin protease was added 1:10 trypsin/sample protein (w/w), centrifuged 1000 rpm for 10 sec, and incubated in dark water bath at 37°C for 13 hours. Peptides were eluted from S-traps with 50 mM ammonium bicarbonate in water, centrifuged at 3,000 rpm for 1 min, repeated elution with 0.1% formic acid in water, and then with 40% acetonitrile containing 0.1% formic acid in water, combining eluates in one 2 mL tube. Peptides were concentrated by evaporation under nitrogen gas stream and reconstituted to a final protein concentration of 1.2 μg/μL in 95% water, 5% acetonitrile, 0.1% formic acid containing 50 nM diluted Pierce^TM^ Peptide Retention Time Calibration Mixture.

Protein digests were analyzed on an UltiMate^TM^ 3000 UHPLC (Thermo Scientific) coupled to an Orbitrap Fusion^TM^ Tribrid mass spectrometer (Thermo Scientific) equipped with EASY-spray^TM^ nano-flow source. Two microgram protein digests in 1 μL injections were loaded onto PepMap^TM^ RSLC C18 NanoSpray column (2 μm, 100Å, 75 μm x 50 cm). Peptides were separated using a solvent gradient with 0.1% formic acid in water (mobile phase A) and 0.1% formic acid in 80% acetonitrile (mobile phase B) at a flow rate of 250 nl min^-1^. For peptide separation, the column was initially equilibrated at 4% B for 3 min., increased to 30% B at 90 min, increased to 55% B at 120 min, increased to 90% B at 130 min, held at 90% B until 134 min., and then decreased to 4% B at 135 min. The solvent gradient included a column flush method that consisted of three rapid gradient flushes of 4% B to 90% B holding at each for 4 min and then re-equilibrated at 4% for 25 min. Peptides were ionized in positive ionization mode using 2.2kV spray voltage, 2 Arb sweep gas flow, and 275°C ion transfer tube temperature. The MS^1^ scan (m/z 300-1,500) was performed in orbitrap mass analyzer at 500,000 resolution with a cycle time of 2 sec. MS^2^ scans were collected for ions that passed the following filters: peptide monoisotopic peak determination, charge states 2-7, dynamic exclusion duration of 40 seconds for 10 ppm mass tolerance, minimum intensity of 1.9E4, and isotope exclusion. MS^2^ scans were acquired in the ion trap mass analyzer with an isolation window of 1.2 amu, following activation with collision-induced dissocation (CID) of 35% energy. Data were processed in Proteome Discoverer (Version 3.1.0.638, Thermo Fisher Scientific) with FDR confidence <0.01, using FASTA files for Mus musculus and Ratus norvegicus: Mus musculus (sp_tr_isoforms_TaxID=10090_and_subtaxon); version 2023-06-28; downloaded on 10/12/2023; 93114 sequences and for Ratus norvegicus (Rattus norvegicus (sp_tr_incl_isoforms TaxID=10116_and_subtaxonomies), downloaded Oct 20; 83181 sequences. Normalized data was used from Proteome discoverer for analysis. Features with>50% missing values were removed, for the remaining missing values were estimated with BPCA. The data was auto scaled.

## Results

### Targeted Chelation Therapy with EDTA NPs acts as Senomorphic and not senolytic

Taking leads from our previous findings and having established that targeted chelation therapy decreases the expression of BMP2and RUNX2, in this study using CKD model, we first validated that EDTA-NP therapy decreases calcium deposition in the aorta and increased longevity in treated animals. We observed a significant decrease in the concentration of inflammatory cytokines, IL-6 and IL-1β in the serum and an improvement in the survival rate of the animals treated with EDTA-NP (Figure 1). *<u>Second</u>*, we compared the transcriptional expression and activity of senescence and SASP markers (SA-β Gal, IL-6, IL-1β, BMP2, MCP1, MMP9, &2) in the aorta harvested from EDTA-NP, Blank NP (BLANK-NP) and control no treatment group animals. We observed a significant decrease in senescence build-up and SASP markers in the aorta harvested from the EDTA-NP treatment group (Figure 2).

**Figure 1.**
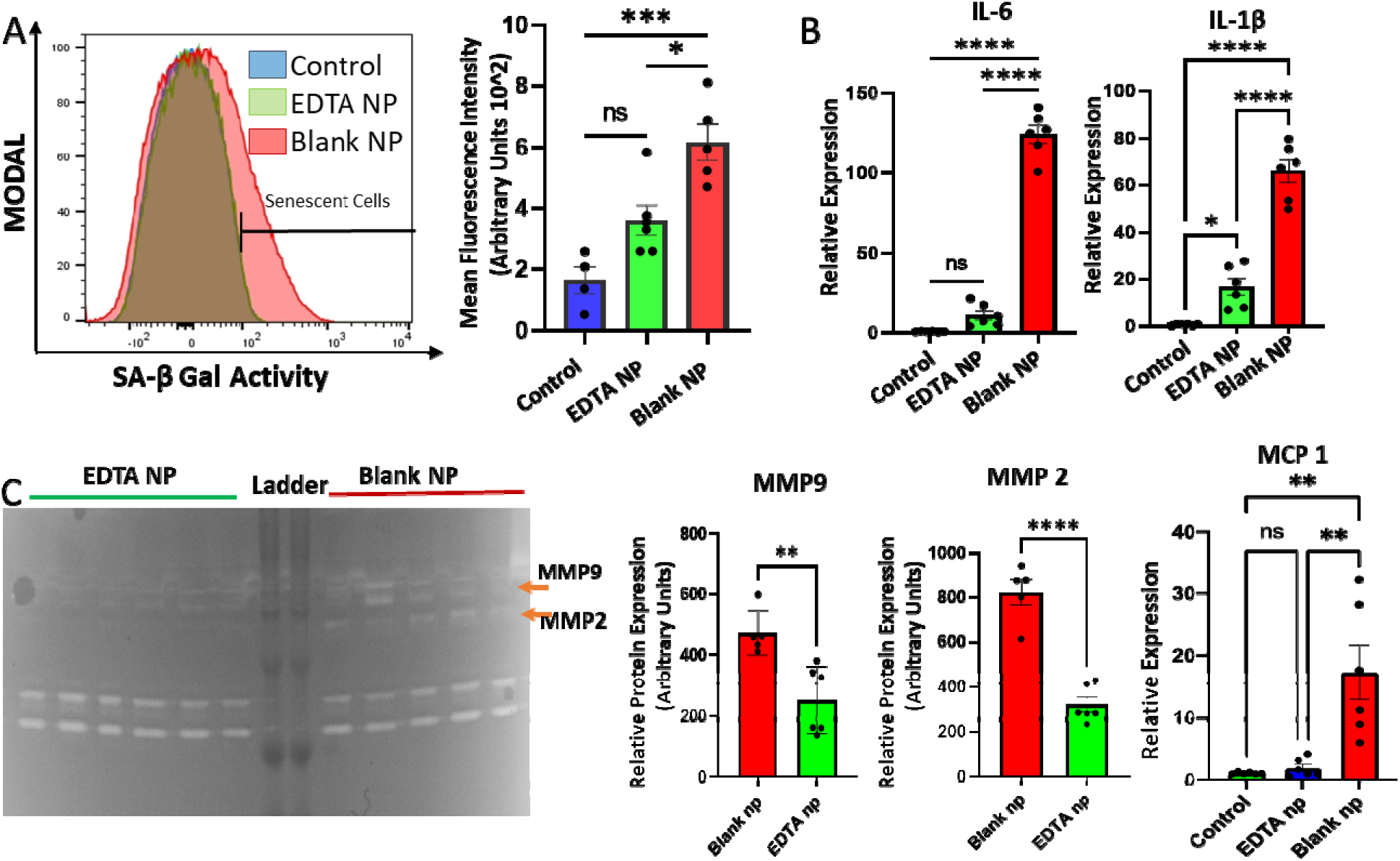
Senomorphic Effect of EDTA nanoparticles. Targeted therapy with EDTA-NP decreased the expression of SASP Markers significantly in aortas harvested from Late-Stage CKD animals- A) SA-β Gal Activity b) transcriptional expression of IL-6, IL-1β, and MCP1 as well as activity of C) MMP 2 & 9. N=6 per group, results represented as Mean±SEM.

**Figure 2.**
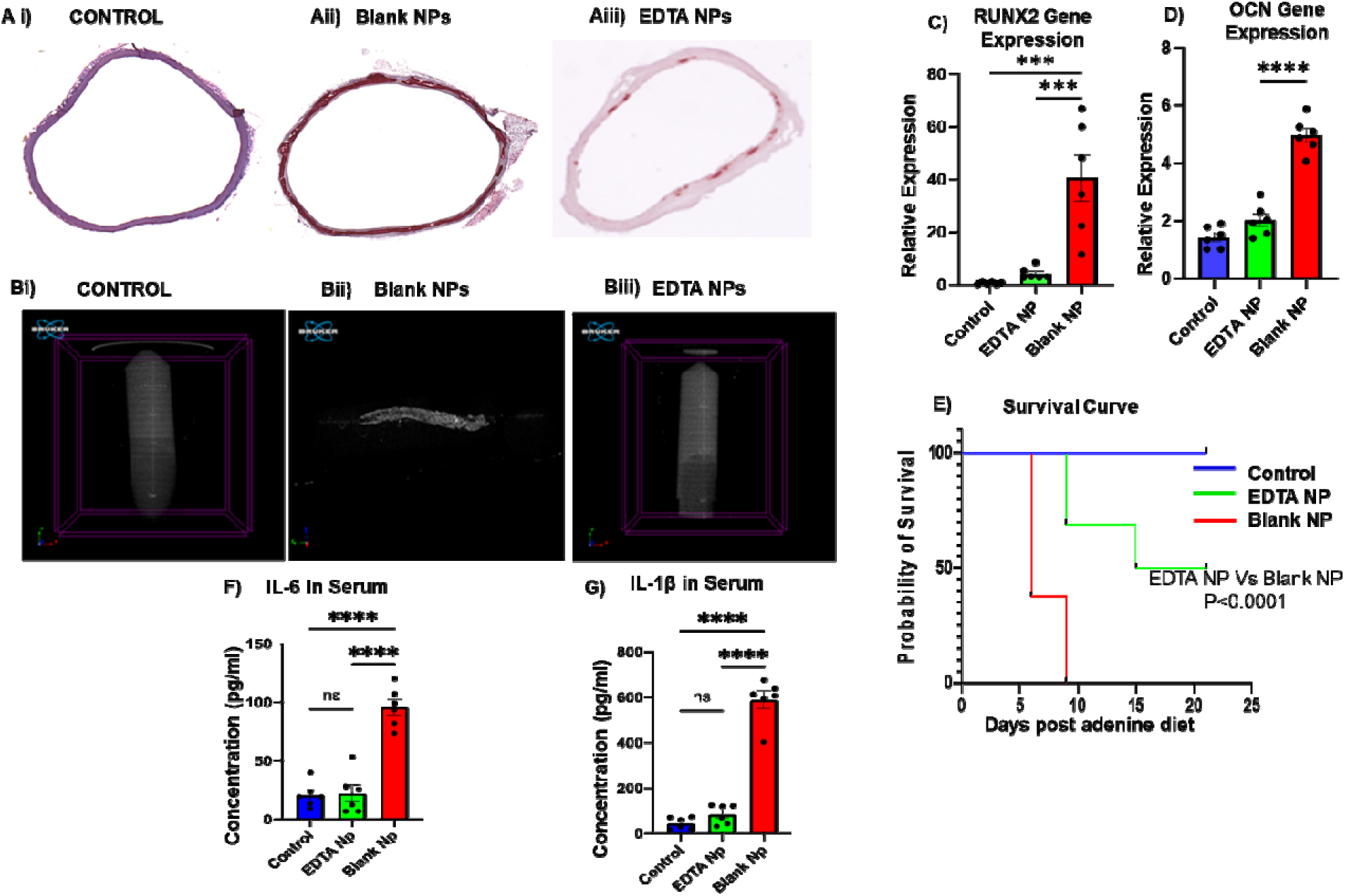
EDTA-NP therapy increases Longevity in Late-Stage CKD rats by decreasing Calcification in Aorta. Calcium deposition in aorta was analyzed by A) Alizarin staining and Micro CT scanning b) Transcript level analysis of ossification markers Osteocalcin (OCN) and RUNX2 and a decrease in their expression level revealed a decreased tendency towards osteoblastic phenotypic switching. Further we observed a significant decrease in circulating levels of pro inflammatory cytokine C) IL-6 and IL-1β with a simultaneous significant improvement in the survival rate as is reflected by the D) Survival curve post adenine diet treatment. N=6 per group results represented as Mean± SEM.

To examine whether EDTA induced apoptosis of senescent cells, we used a well-established senolytic agent ABT 263, which is known to induce apoptosis by recruiting NLRP3 via caspase3 activation and tested it against EDTA. We observed that ABT 263 treatment increased caspase-3 expression, whereas EDTA treatment instead decreased Caspase 3 expression in primary human vascular smooth muscle cells exposed to high phosphate ion concentration, indicating that EDTA does not induce NLRP3 mediated apoptosis. Furthermore, a decrease in Caspase 3 expression suggests transcriptional inhibition of NLRP3 expression during the priming stage (Figure 3 A, B)

**Figure 3A:**
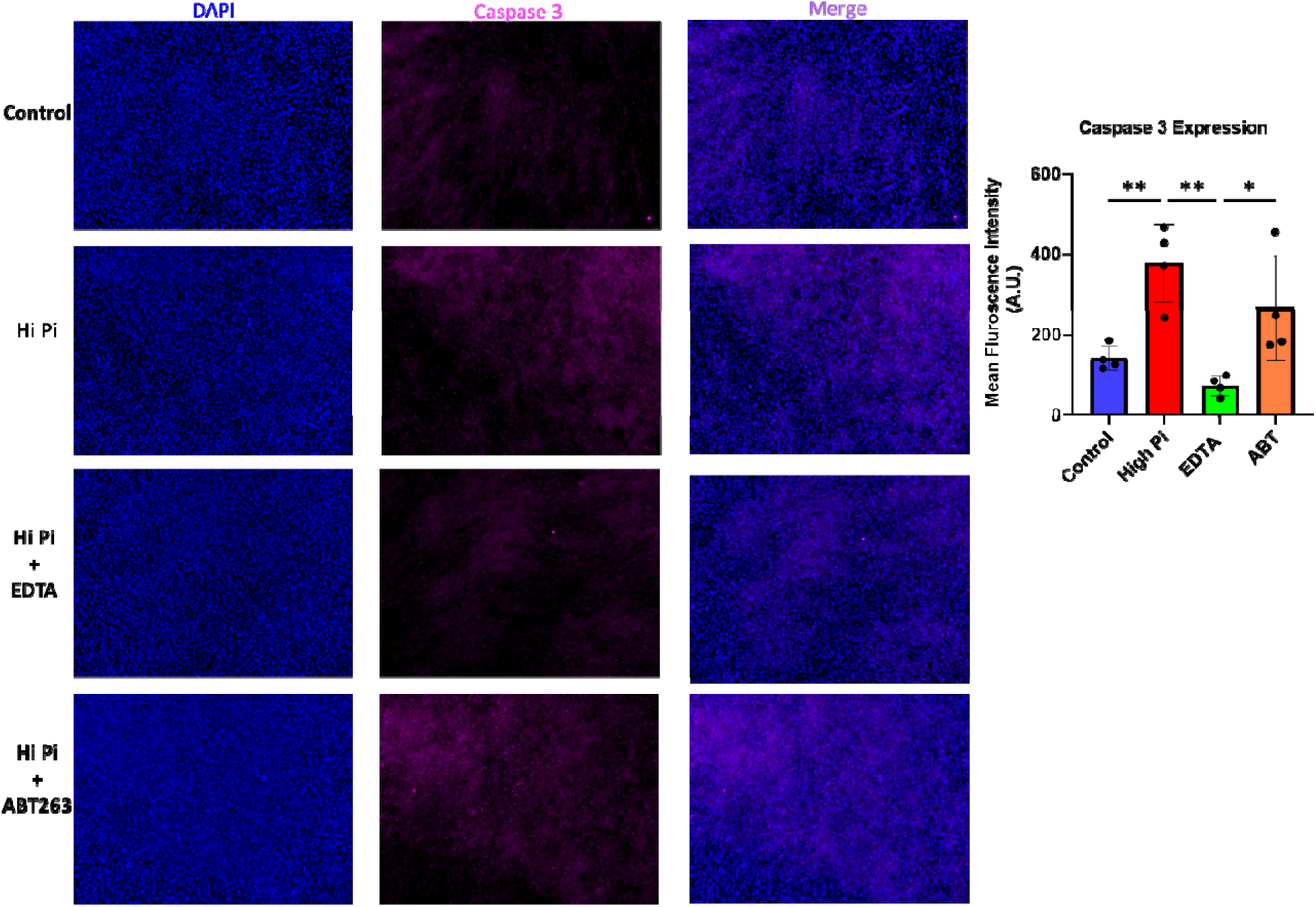
EDTA is Senomorphic and not Senolytic. Immortalized human vascular smooth muscle cells cultured under high phosphate conditions for 3 days and were treated with either EDTA or ABT 263 (senolytic agent) for 24 hours. IHC based Expression level analysis of Caspase 3 was used as marker for senolytic (inducing selective apoptosis of senescent cells) activity. We observed that in comparison to ABT 263, EDTA treatment significantly brought down Caspase 3 activity, indicating that EDTA does not induce apoptosis. Experiment performed in triplicate repeated 4 times, results represented as Mean±S.D.

**Figure 3b:**
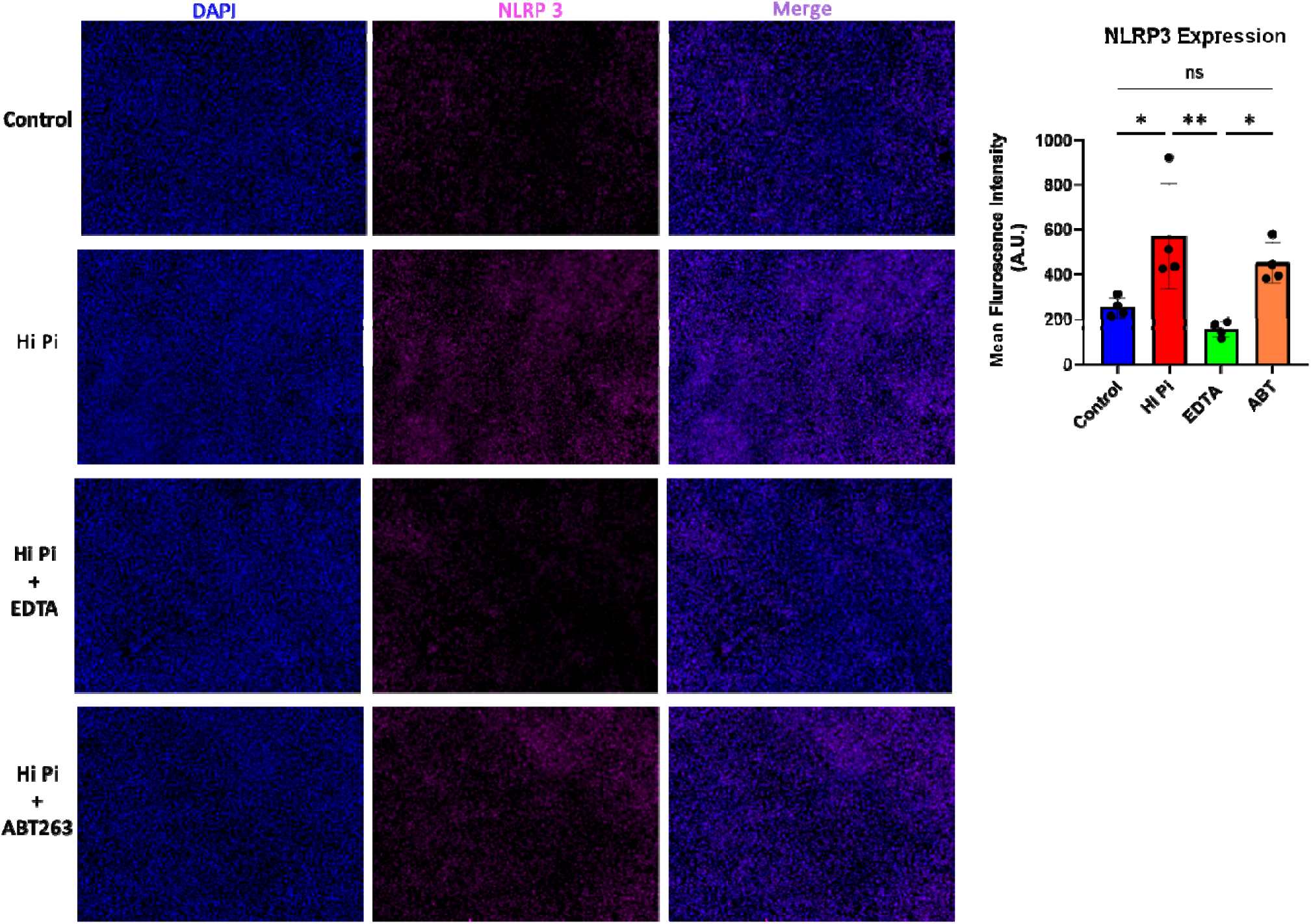
EDTA but not ABT263 Decrease NLRP3 expression. Immortalized human vascular smooth muscle cells cultured under high phosphate conditions for 3 days and were treated with either EDTA or ABT 263 (senolytic agent) for 24 hours. IHC based Expression level analysis of NLRP 3 reveals that EDTA treatment and not ABT 263 decreased NLRP 3 activity. Justifying anti-apoptotic and senomorphic nature of EDTA treatment. Experiment performed in triplicate repeated 4 times, results represented as Mean±S.D.

### Targeted chelation therapy with EDTA NPs Significantly decreased NLRP3 Expression in Calcified Aorta

We examined the expression of NLRP3, in the calcified aorta harvested from late-stage CKD rats as well as in the long-duration aortic ring culture model. In the calcified aortas harvested from both aortic ring culture and CKD model, we observed that treatment with EDTA and EDTA-NP, respectively, caused a significant decrease in NLRP3 expression as quantified by IHC, and qPCR as well as a decrease in the concentration of IL-1 β and IL-6 in the culture supernatants (Figure 4 A, B).

**Figure 4.**
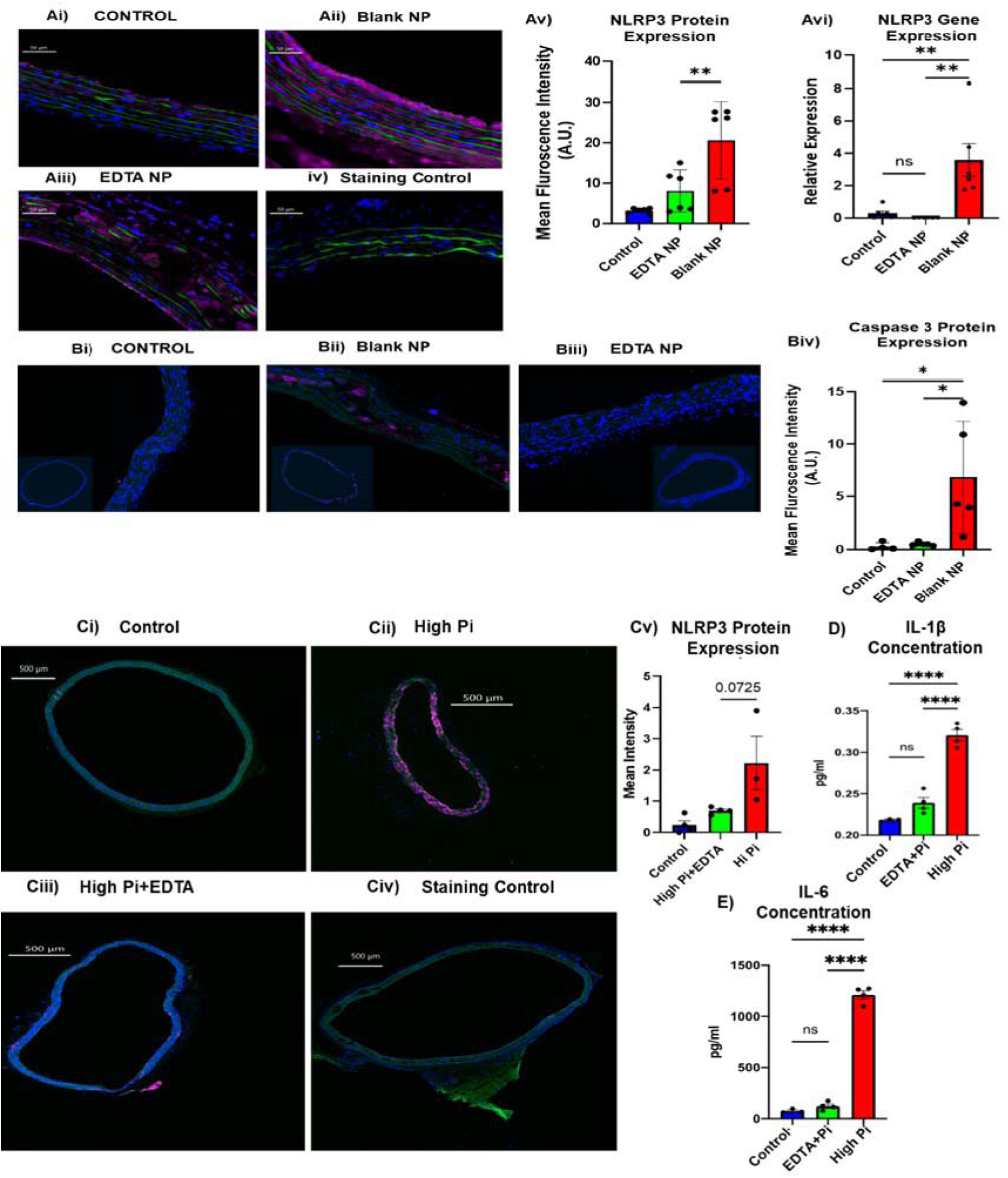
Effect of EDTA-NP treatment on NLRP3 and Caspase3 expression in A-B) calcified aortas harvested from late-stage in vivo CKD model- (A separate cohort of N=5 per group was set up, results represented as Mean ± SEM); **C)** calcified aortas harvested after 10 days of ex-vivo/Long-term culture under high phosphate conditions (N=4 per group; experiment done in triplicate; results represented as Mean± SD). EDTA treatment significantly decreased Caspase 3 and NLRP3 expression in the late-stage in-vivo CKD model of vascular calcification as well as in the long term in-vitro model

### Proteomics Analysis reveals a significant decrease in SASP markers and simultaneous increase in markers of Angiogenesis

We performed an untargeted top-down proteomics analysis of total protein isolated from abdominal aorta of EDTA-NP treated Vs Blank-NP treated animal. The analysis revealed that 498 proteins were exclusively expressed in Blank-NP group and 200 proteins were exclusively expressed in the EDTA-NP treatment group (Figure 5A). A total of 825 proteins were identified that were significantly differentially expressed between two groups (Figure 5B, Supp. Table 2). Amongst the top 25 upregulated proteins, 26% are directly influence cell proliferation and angiogenesis (Anaxa8, Enpep, Fgf8b,Clec11a, Crlf1, Agc1, Lancl2), 14% prevent osteoblastic transition of VSMCs (MGP, Phosphorylated form of Spp1, and ANKH), 14% play a role in cytoskeleton stabilization (RCG23467, Dmd, and Actb), 10% are involved in mediating proper protein folding (Ppidl1, Hmga1), 10% are involved in anabolic Lipid and carbohydrate metabolism (Mdh1 Mor2, Pla2g4a), and another 10 and 5% are anticoagulant and coagulants (Proc, and F7) (Figure 5C).

**Figure 5.**
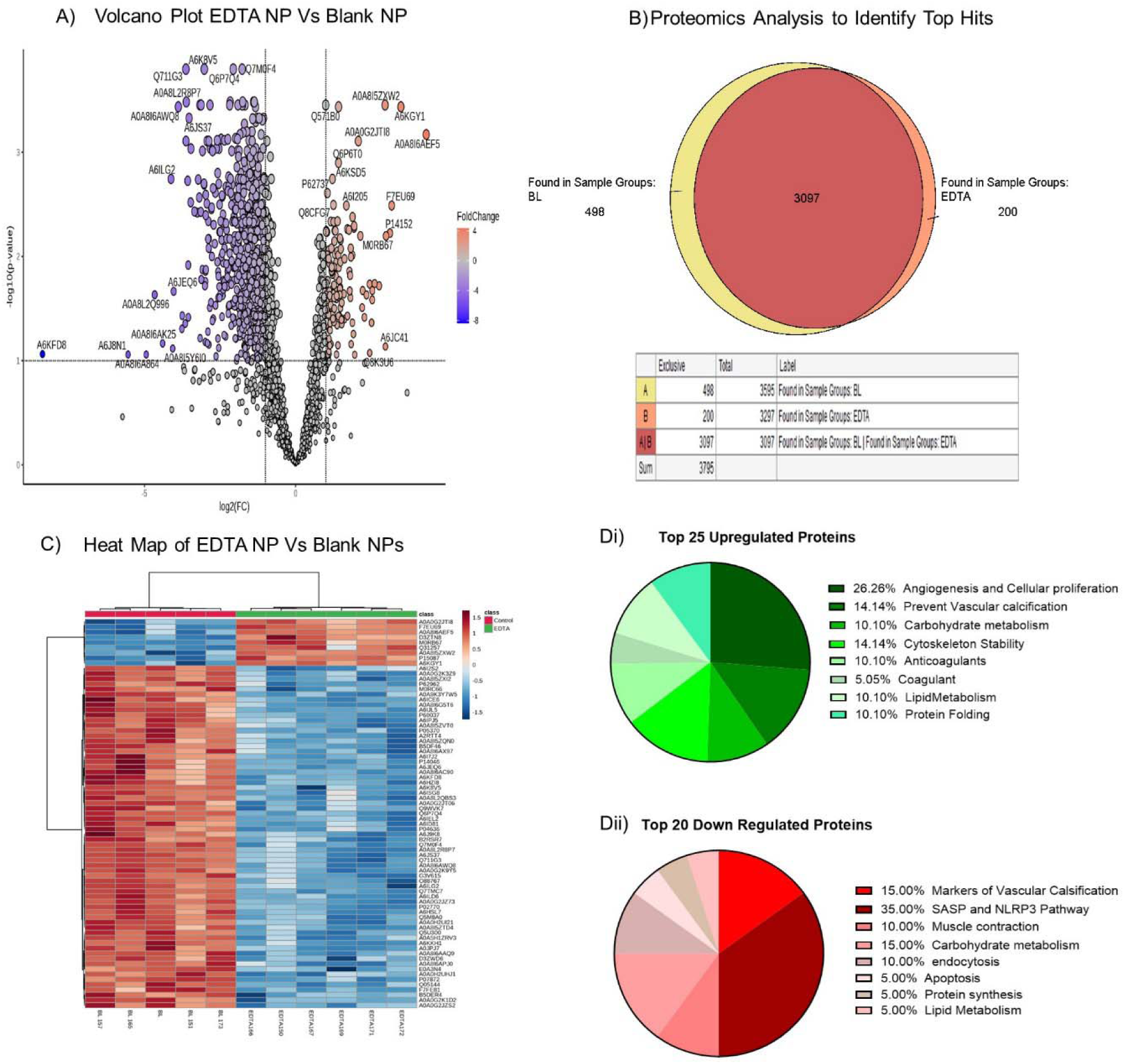
Proteomic Analysis of abdominal aorta from EDTA NP treated group and Blank NP treated group. **A)** Breakdown of total proteins identified during the analysis as exclusively expressed or differentially expressed in both the treatment groups. **B)** A volcano plot for the significantly differentially expressed proteins-Upregulated (purple) or downregulated (red) in EDTA treatment Vs Blank treatment group. C) Heat Map of differentially Expressed proteins found in all the samples **Di-ii)** Breakdown of top 25 upregulated and top 20 downregulated proteins respectively as per biological functions. N≥5 per group.

Amongst the top 20 downregulated proteins, 35% were directly involved in facilitating SASP processes (Uchl1, H1-1 Hist1h1a, HP, Cfb C2, Tf, Serpina1), 15% were markers of vascular calcification (Ckmt2, Fetub, Pvalb Pva), 10% regulate muscle contraction (Myl2, RCG36716), 5% are involved in each Lipid metabolism, protein synthesis and apoptosis (Iah1 Harpb64, Eef1a2 Kcnq2, Kng1) (Figure 5D). We screened the list of downregulated proteins to scrutinize the ones that regulate NLRP3 activation. Csnk2b, Rbp4 regulate phase 1 of NLRP3 activation via Jak2/MapK/P13/AK. Protein S100a9 and Tf initiate phase 2 of NLRP3 activation and Jpt1 Sumo2, and PyCad are involved in stabilization of NLRP3 inflammasome complex. Amongst other downregulated proteins were the markers of senescence and SASP-Smap1, Serpina3k Serpin2b, Bcl2l13, Bin1.

## Discussion

The presence of senescent cells plays a vital role in the progression of pathology (like diabetes and chronic kidney disorder) as well as aging-associated vascular calcification. In context to vascular calcification observed due to chronic kidney disorder, mineral imbalance, and high intracellular concentration of Ca and P ions, which develop in patients with CKD stage 4 or 5, is critical for precipitating phenotypic changes in tunica media (32–34). Clinical data from CKD patients also shows the presence of SASP and senescent cells in the kidney that coincide with the appearance of medial artery calcification (35,36). Even though senescent cells are in the stage of cell cycle arrest, these cells are metabolically active and possess a distinct inflammatory secretory phenotype (Senescence associated secretory phenotype, SASP) that has the potential to modulate tissue microenvironment and in precipitating phenotypic transitions in the neighboring healthy cells within the tissue (30,37). Significant number of in-vitro reports highlight the potential role of the SASP in osteoblastic differentiation of VSMCs. However, both the source of SASP and the translation of senolytic/senomorphics to clinics for treating cardiovascular diseases are still a subject of debate (38,39). Despite promising preclinical studies, clinical translation of senolytics for treating diseases is hindered mainly because of their off-targeting side effects and excessive apoptosis-induced hemorrhages. A recent study by Grosse et al (39) demonstrated that the global elimination of P16^high^ cells was non-replaceable and was, in fact, detrimental to health and lifespan as it led to the liver and cardiac fibrosis as well as disruption of the blood tissue barrier. This emphasizes the need to investigate the local source of SASP further and harness a targeted drug delivery approach to modulate the vascular microenvironment for treating senescence-associated vascular calcification encountered during CKD. Recently NLRP3 inflammasome activation in the calcified and inflammatory cardiovascular cells has emerged as an important source of SASP and a biomarker of cellular senescence. Our lab has also previously shown that culturing VSMCs under hyperphosphatemia conditions induces their transformation into osteoblast-like cells and upregulates the expression of biomarkers of cellular senescence specifically BMP2 and RUMX2 in the VSMCs (21). We have also previously established that targeted chelation therapy with EDTA-NP can cause a reversal of vascular calcification and reverses the phenotype of ossified VSMCs in the aorta (20).

Therefore, in the present study we investigated whether targeted chelation therapy with EDTA-NP has serotherapeutic potential. We used a previously established flow cytometry-based assay to detect the presence of senescent cells from the primary cells harvested from the tissue immediately after tissue harvesting. This method works because senescent cells express a relatively higher SA-β Gal activity-gold standard marker for senescence. Using this standardized assay to measure SA-βGal activity, we were able to evidence a decrease in percentage of senescent cells in the aortas harvested from the EDTA NP treatment group in comparison to the Blank-NP treated animals.

Next, we wanted to address whether the decrease in the senescent cell accumulation observed was due to senolytic (induction of selective apoptosis of senescent cells) property of EDTA. For this purpose, we compared EDTA with a well-established senolytic agent ABT263. ABT263 activates Caspase3 pathway by recruiting NLRP3 and induces apoptosis selectively in senescent cells (40). In our study we observed that unlike ABT263; EDTA treatment decreased Caspase3 and NLRP3 expression in the VSMCs maintained under high Pi condition whereas ABT263 had no impact either on NLRP3 activity. Even though we expected ABT to show an increase in NLRP3 and Caspase 1 expression but a deviation from our expected results might have resulted because the IHC expression of these proteins was not normalized against cell count.

Next we evaluated the effect of EDTA chelation therapy on the NLRP3, Caspase1, and IL-1β expression in the abdominal aorta samples harvested from the CKD animal model and we observed a similar trend, a decrease in NLRP3, Caspase1, IL-1β expression in the EDTA-NP vs Blank-NP treatment group.

Recent research documents the role of NLRP3 inflammasome activation as a source of SASP. Its activation has been linked to precipitation of senescent phenotype in chondrocytes, osteoblasts, and adipocytes. NLRP3 activation has also been evidenced to promote arterial calcification by causing pyroptosis (13–16). Its activation leads to the conversion of pro-caspase1 to caspase 1, ultimately inducing cell lysis. The overload of apoptotic debris produced in the process serves as the scaffold for Ca and P deposition and causes the remodeling of arterial tissue. In addition to its involvement in direct tissue remodeling, NLRP3 inflammasome activation also activates innate immune machinery in cells; it triggers the maturation of proinflammatory cytokines such as interleukin-1beta to engage innate immune defenses and hence plays a critical role in the progression of disease. In context to chronic kidney disorder and aging-associated vascular calcification, experimental data show that NLRP3 inflammasome pathway is activated in kidney and VSMCs by klotho deficiency and doxorubicin (a well-known senescence inducer) (15,41).

In this study, we observed that EDTA chelation therapy decreases PIT-1 expression in calcified aorta and proteomic analysis revealed significant downregulation in S100A9 and Signal recognition particle receptor both of which translate intracellular calcium concentration to activate NLRP3 inflammasome assembly. Furthermore, we observed a significant downregulation of PYD and CARD domain containing peptide, which is primarily responsible for recruitment of proCaspase1, and its conversion to Caspase 1 by NLRP3 inflammasome complex. We also observed that the EDTA-NP therapy decreased the expression of proteins that regulate osteogenic differentiation of VSMCs ( Ckm2, FeutB, Vitamin D-binding protein) and increased the expression of proteins involved in angiogenesis (Enpep, Osteopontin and MGP) in the aortas harvested from the chronic kidney disease-based animal model of MAC. The animals also survived longer than the control group, and we observed a significant decrease in SASP markers -Serpin1, serpin 3, BCL2, BIN. Our findings serve as proof of concept for using agents that can modulate ionic imbalance in tissue microenvironment as senomorphics for targeting senescent cells to treat vascular stenosis by creating conditions suitable for vascular cells to reprogram their phenotype.

## Limitations

The current study was performed to evaluate the role of a high phosphate environment on initiating senescence phenotype in aortic tissue overall. A question how mineral imbalance turns on SASP intracellularly remains unanswered. Even though the upregulation of PiT-1 expression hints towards the involvement of Pi^+^ ions, experimental evidence is still required to establish a link. The other interesting aspect is to investigate the pathways responsible for senomorphic activity of EDTA. Further, we acknowledge that the study would have benefited from the use of a direct pharmacological inhibitor of NLRP3.

## Supporting information

Early-Stage vascular calcification in Chronic Kidney Disorder (CKD) Model

Aortic Ring Culture for vascular calcification

A list of Primers used (OCN, RUNX2, IL-6, IL-1β, MCP-1, p21, p19, and GAPDH) in the study is given in supplementary Table-1.

## Acknowledgements

This research was funded partially by grant funding from National Institutes of Health to NRV (R01HL133662, R01HL145064, P30GM131959). We would like to thank Dr. Agnes Nagy-Mehesz at S.C. Bio Craft COBRE Core and Dr. Nishanth Tharayil as well as Elizabeth Leonard at the MUAL - Multi User analytical lab and Metabolomics Core, at Clemson for their help and support.

## Data availability statement

All the relevant data is provided in the manuscript. The data that support the findings of this study are available from the corresponding authors, [NRV and SA], upon reasonable request.

## Supplementary Table 1

**Table 1.**
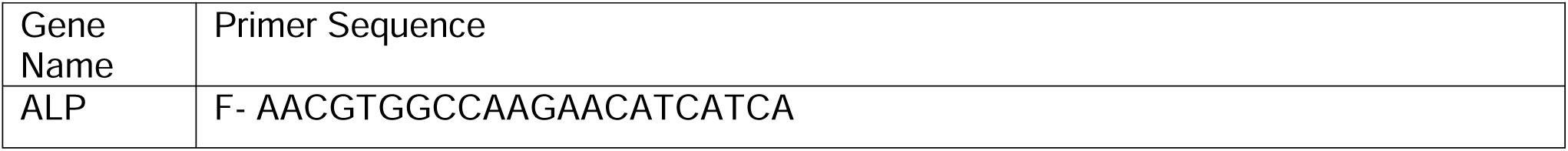
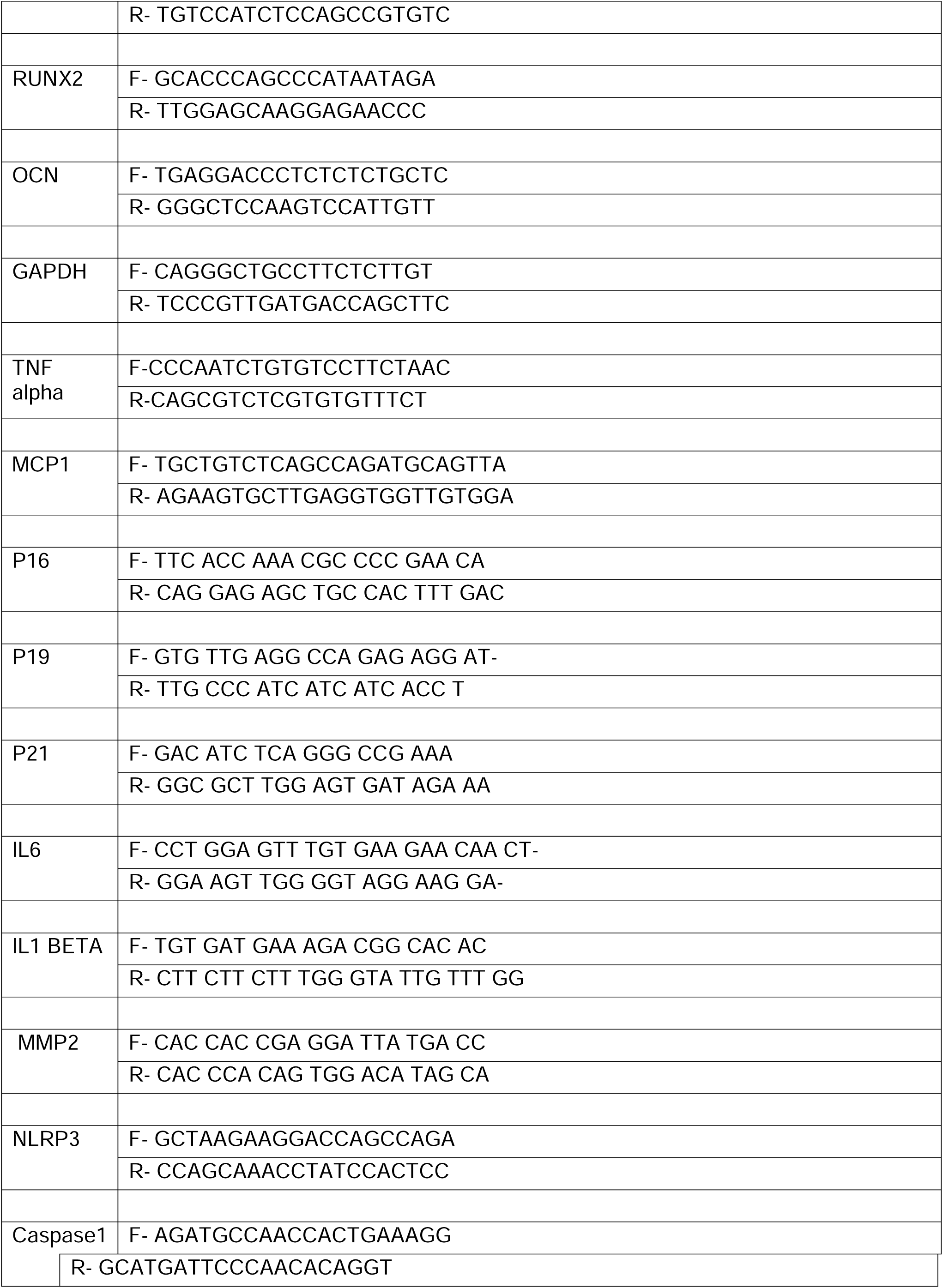
Primer Sequence.

## References

1. Dai L, Qureshi AR, Witasp A, Lindholm B, Stenvinkel P. Early Vascular Ageing and Cellular Senescence in Chronic Kidney Disease. Comput Struct Biotechnol J. 2019 Jan 1;17:721–9.

2. Schlieper G, Schurgers L, Brandenburg V, Reutelingsperger C, Floege J. Vascular calcification in chronic kidney disease: an update. Nephrol Dial Transplant [Internet]. 2016 [cited 2022 Dec 19];31:31–9. Available from: https://academic.oup.com/ndt/article/31/1/31/2459964

3. Fang YP, Zhao Y, Huang JY, Yang X, Liu Y, Zhang XL. The functional role of cellular senescence during vascular calcification in chronic kidney disease. Front Endocrinol (Lausanne). 2024 Jan 22;15.

4. Oh KS, Febres-Aldana CA, Kuritzky N, Ujueta F, Arenas IA, Sriganeshan V, et al. Cellular senescence evaluated by P16INK4a immunohistochemistry is a prevalent phenomenon in advanced calcific aortic valve disease. Cardiovascular Pathology. 2021 May;52:107318.

5. Roos CM, Zhang B, Palmer AK, Ogrodnik MB, Pirtskhalava T, Thalji NM, et al. Chronic senolytic treatment alleviates established vasomotor dysfunction in aged or atherosclerotic mice. Aging Cell. 2016 Oct 5;15(5):973–7.

6. Bennett G. CHILDS DJBTWCACJCAJMVD. Senescent intimal foam cells are deleterious at all stages of atherosclerosis. 2016.

7. Childs BG, Gluscevic M, Baker DJ, Laberge RM, Marquess D, Dananberg J, et al. Senescent cells: an emerging target for diseases of ageing. 2017 [cited 2023 Jan 26]; Available from: www.nature.com/nrd

8. Nguyen NT, Nguyen TT, Da Ly D, Xia JB, Qi XF, Lee IK, et al. Oxidative stress by Ca overload is critical for phosphate-induced vascular calcification. American Journal of Physiology-Heart and Circulatory Physiology. 2020 Dec 1;319(6):H1302–12.

9. Melinda Duer AMCCMS. DNA Damage Response A Molecular Lynchpin in the Pathobiology of Arteriosclerotic Calcification. Arterioscler Thromb Vasc Biol. 2020;e193–202.

10. Nakano-Kurimoto R, Ikeda K, Uraoka M, Nakagawa Y, Yutaka K, Koide M, et al. Replicative senescence of vascular smooth muscle cells enhances the calcification through initiating the osteoblastic transition. American Journal of Physiology-Heart and Circulatory Physiology. 2009 Nov;297(5):H1673–84.

11. Zuccolo E, Badi I, Scavello F, Gambuzza I, Mancinelli L, Macrì F, et al. The microRNA-34a-Induced Senescence-Associated Secretory Phenotype (SASP) Favors Vascular Smooth Muscle Cells Calcification. Int J Mol Sci. 2020 Jun 23;21(12):4454.

12. Burton DGA, Matsubara H, Ikeda K. Pathophysiology of vascular calcification: Pivotal role of cellular senescence in vascular smooth muscle cells. Exp Gerontol. 2010 Nov 1;45(11):819–24.

13. Liao LZ, Chen ZC, Wang SS, Liu WB, Zhao CL, Zhuang XD. NLRP3 inflammasome activation contributes to the pathogenesis of cardiocytes aging. 2021 [cited 2023 Jan 2];13(16). Available from: www.aging-us.com

14. Pinar AA, Scott TE, Huuskes BM, Tapia Cáceres FE, Kemp-Harper BK, Samuel CS. Targeting the NLRP3 inflammasome to treat cardiovascular fibrosis. 2020 [cited 2023 Oct 15]; Available from: 10.1016/j.pharmthera.2020.107511

15. Li X, Li Z, Li B, Zhu X, Lai X. Klotho improves diabetic cardiomyopathy by suppressing the NLRP3 inflammasome pathway. Life Sci. 2019 Oct;234:116773.

16. Haneklaus M, O’Neill LAJ. NLRP3 at the interface of metabolism and inflammation. Immunol Rev. 2015 May;265(1):53–62.

17. Poudel B, Gurung P. An update on cell intrinsic negative regulators of the NLRP3 inflammasome. J Leukoc Biol. 2018 Jun;103(6):1165–77.

18. Yu C, Zhang C, Kuang Z, Zheng Q. The Role of NLRP3 Inflammasome Activities in Bone Diseases and Vascular Calcification. Inflammation. 2021 Apr 20;44(2):434–49.

19. Ho LC, Chen YH, Wu TY, Kao LZ, Hung SY, Liou HH, et al. Phosphate burden induces vascular calcification through a NLRP3-caspase-1-mediated pyroptotic pathway. Life Sci. 2023 Nov;332:122123.

20. Zohora FT, Arora S, Swiss A, Vyavahare N. Reversal of heavy arterial calcification in a rat model of chronic kidney disease using targeted ethylene diamine tetraacetic acid-loaded albumin nanoparticles. Cardiovasc Diagn Ther. 2024 Aug;14(4):489–508.

21. Lei Y, Nosoudi N, Vyavahare N. Targeted chelation therapy with EDTA-loaded albumin nanoparticles regresses arterial calcification without causing systemic side effects. Journal of Controlled Release. 2014 Dec;196:79–86.

22. Kaneda A, Fujita T, Anai M, Yamamoto S, Nagae G, Morikawa M, et al. Activation of Bmp2-Smad1 Signal and Its Regulation by Coordinated Alteration of H3K27 Trimethylation in Ras-Induced Senescence. PLoS Genet. 2011 Nov 3;7(11):e1002359.

23. Camell CD, Günther P, Lee A, Goldberg EL, Spadaro O, Youm YH, et al. Aging Induces an Nlrp3 Inflammasome-Dependent Expansion of Adipose B Cells That Impairs Metabolic Homeostasis. Cell Metab. 2019 Dec 3;30(6):1024–1039.e6.

24. Ni B, Pei W, Qu Y, Zhang R, Chu X, Wang Y, et al. MCC950, the NLRP3 Inhibitor, Protects against Cartilage Degradation in a Mouse Model of Osteoarthritis. Oxid Med Cell Longev. 2021 Nov 3;2021:1–14.

25. Cordero MD, Williams MR, Ryffel B. AMP-Activated Protein Kinase Regulation of the NLRP3 Inflammasome during Aging. Trends in Endocrinology and Metabolism. 2018 Jan 1;29(1):8–17.

26. Sinha A, Shaporev A, Nosoudi N, Lei Y, Vertegel A, Lessner S, et al. Nanoparticle targeting to diseased vasculature for imaging and therapy. Nanomedicine. 2014 Jul;10(5):e1003–12.

27. Nosoudi N, Nahar-Gohad P, Sinha A, Chowdhury A, Gerard P, Carsten CG, et al. Prevention of Abdominal Aortic Aneurysm Progression by Targeted Inhibition of Matrix Metalloproteinase Activity With Batimastat-Loaded Nanoparticles. Circ Res. 2015 Nov 6;117(11).

28. Zohora FT, Arora S, Swiss A, Vyavahare N. Reversal of heavy arterial calcification in a rat model of chronic kidney disease using targeted ethylene diamine tetraacetic acid-loaded albumin nanoparticles. Cardiovasc Diagn Ther. 2024 Aug;14(4):489–508.

29. Vyavahare N and Sinha A. Formation of Delivery Agents targeted to degraded elastic fibers. US Patent, 2020, Patent number 10,688,061.

30. Arora S, Thompson PJ, Wang Y, Bhattacharyya A, Apostolopoulou H, Hatano R, et al. Invariant natural killer T cells coordinate removal of senescent cells. Med. 2021 Aug;2(8):938–950.e8.

31. Arora S, Vyavahare N. Elastin-targeted nanoparticles delivering doxycycline mitigate cytokine storm and reduce immune cell infiltration in LPS-mediated lung inflammation. PLoS One. 2023 Jun 1;18(6 June).

32. Hutcheson JD, Goettsch C. Cardiovascular Calcification Heterogeneity in Chronic Kidney Disease. Circ Res. 2023 Apr 14;132(8):993–1012.

33. Palit S, Kendrick J. Vascular Calcification in Chronic Kidney Disease: Role of Disordered Mineral Metabolism. Curr Pharm Des. 2014 Feb 12;20(37):5829–33.

34. Dube P, DeRiso A, Patel M, Battepati D, Khatib-Shahidi B, Sharma H, et al. Vascular Calcification in Chronic Kidney Disease: Diversity in the Vessel Wall. Biomedicines. 2021 Apr 8;9(4):404.

35. Yamada S, Tokumoto M, Tatsumoto N, Tsuruya K, Kitazono T, Ooboshi H. Very low protein diet enhances inflammation, malnutrition, and vascular calcification in uremic rats. Life Sci. 2016 Feb 1;146:117–23.

36. Price PA, Roublick AM, Williamson MK. Artery calcification in uremic rats is increased by a low protein diet and prevented by treatment with ibandronate. Kidney Int [Internet]. 2006 Nov [cited 2023 Dec 5];70(9):1577–83. Available from: https://pubmed.ncbi.nlm.nih.gov/16955099/

37. Thompson PJ, Shah A, Ntranos V, Van Gool F, Atkinson M, Bhushan A. Targeted Elimination of Senescent Beta Cells Prevents Type 1 Diabetes. Cell Metab. 2019 May 7;29(5):1045–1060.e10.

38. Owens WA, Walaszczyk A, Spyridopoulos I, Dookun E, Richardson GD. Senescence and senolytics in cardiovascular disease: Promise and potential pitfalls. Mech Ageing Dev. 2021 Sep;198:111540.

39. Grosse L, Wagner N, Emelyanov A, Molina C, Lacas-Gervais S, Wagner KD, et al. Defined p16High Senescent Cell Types Are Indispensable for Mouse Healthspan. Cell Metab. 2020 Jul;32(1):87–99.e6.

40. Hu L, Chen M, Chen X, Zhao C, Fang Z, Wang H, et al. Chemotherapy-induced pyroptosis is mediated by BAK/BAX-caspase-3-GSDME pathway and inhibited by 2-bromopalmitate. [cited 2023 Apr 12]; Available from: 10.1038/s41419-020-2476-2

41. Xiang T, Luo X, Ye L, Huang H, Wu Y. Klotho alleviates NLRP3 inflammasome-mediated neuroinflammation in a temporal lobe epilepsy rat model by activating the Nrf2 signaling pathway. Epilepsy & Behavior. 2022 Mar;128:108509.

