## Supplementary figures and images for "Targeted chelation therapy decreases NLRP3 expression by vascular cells and acts as senomorphic in Chronic Kidney Disorder induced Vascular Calcification"

### Aortic Ring Culture for vascular calcification

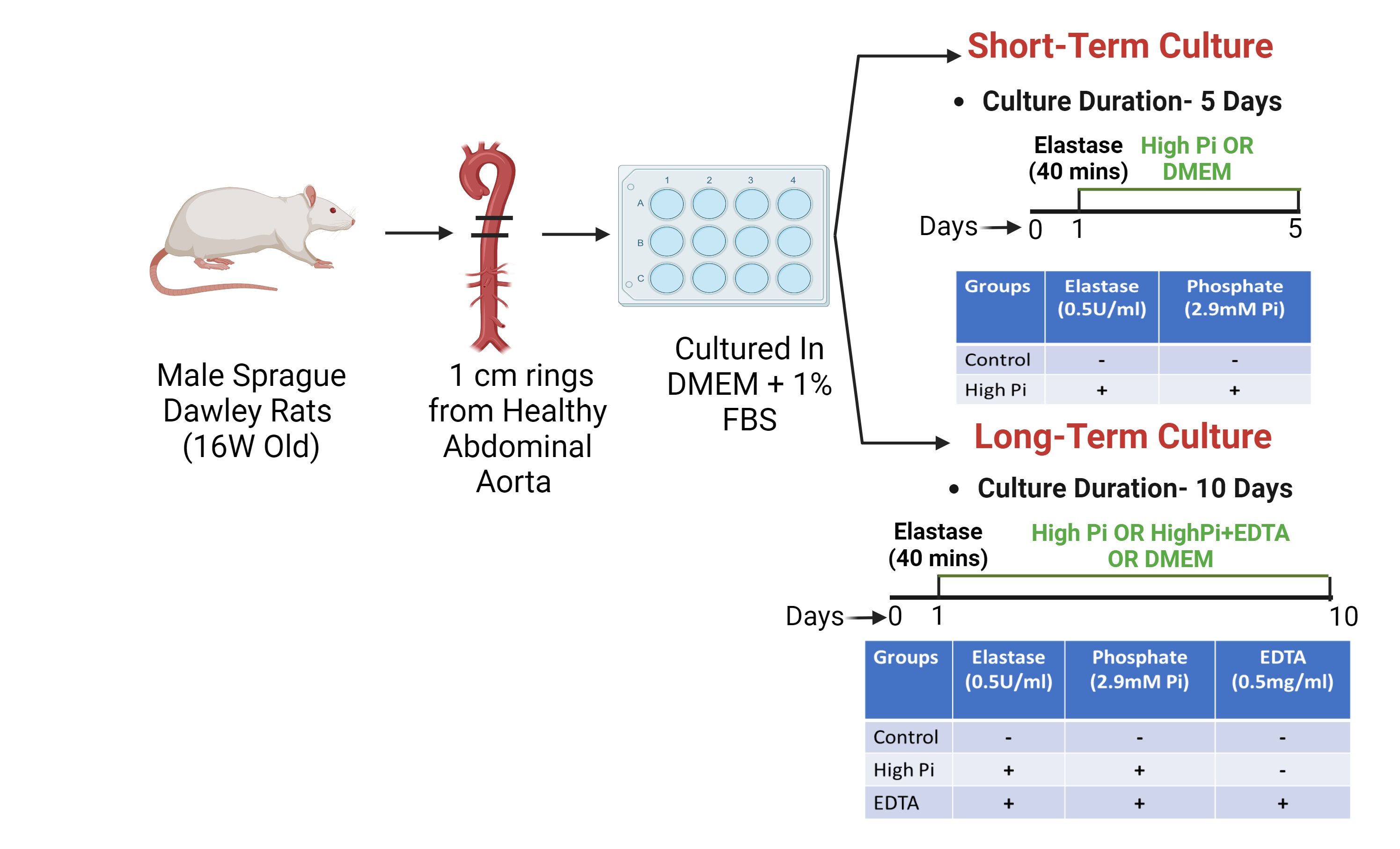

### Early-Stage vascular calcification in Chronic Kidney Disorder (CKD) Model

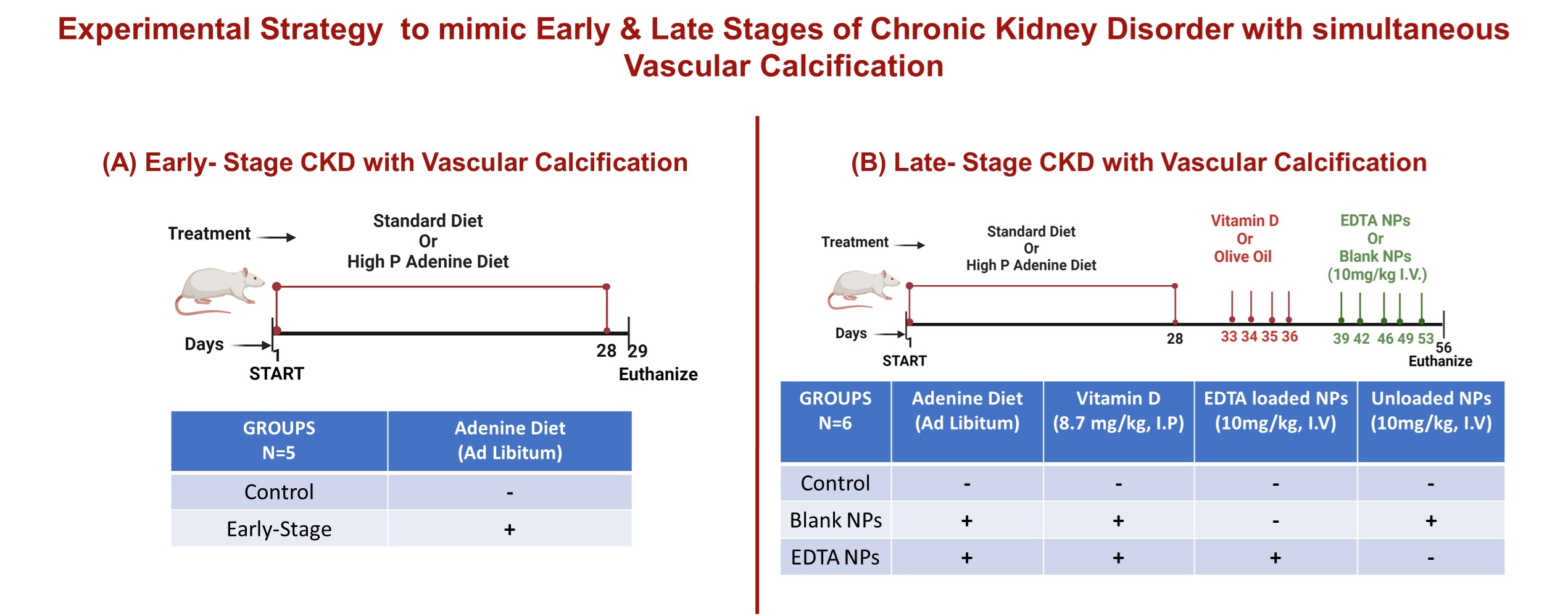
