## Supplementary material for "Targeted chelation therapy decreases NLRP3 expression by vascular cells and acts as senomorphic in Chronic Kidney Disorder induced Vascular Calcification": A list of Primers used (OCN, RUNX2, IL-6, IL-1β, MCP-1, p21, p19, and GAPDH) in the study is given in supplementary Table-1.

Table 1- Primer Sequence

| Gene Name | Primer Sequence |
| --- | --- |
| ALP | F- AACGTGGCCAAGAACATCATCA |
|  | R- TGTCCATCTCCAGCCGTGTC |
| RUNX2 | F- GCACCCAGCCCATAATAGA |
|  | R- TTGGAGCAAGGAGAACCC |
| OCN | F- TGAGGACCCTCTCTCTGCTC |
|  | R- GGGCTCCAAGTCCATTGTT |
| GAPDH | F- CAGGGCTGCCTTCTCTTGT |
|  | R- TCCCGTTGATGACCAGCTTC |
| TNF alpha | F-CCCAATCTGTGTCCTTCTAAC |
|  | R-CAGCGTCTCGTGTGTTTCT |
| MCP1 | F- TGCTGTCTCAGCCAGATGCAGTTA |
|  | R- AGAAGTGCTTGAGGTGGTTGTGGA |
| P16 | F- TTC ACC AAA CGC CCC GAA CA |
|  | R- CAG GAG AGC TGC CAC TTT GAC |
| P19 | F- GTG TTG AGG CCA GAG AGG AT- |
|  | R- TTG CCC ATC ATC ATC ACC T |
| P21 | F- GAC ATC TCA GGG CCG AAA |
|  | R- GGC GCT TGG AGT GAT AGA AA |
| IL6 | F- CCT GGA GTT TGT GAA GAA CAA CT- |
|  | R- GGA AGT TGG GGT AGG AAG GA- |
| IL1 BETA | F- TGT GAT GAA AGA CGG CAC AC |
|  | R- CTT CTT CTT TGG GTA TTG TTT GG |
| MMP2 | F- CAC CAC CGA GGA TTA TGA CC |
|  | R- CAC CCA CAG TGG ACA TAG CA |
| NLRP3 | F- GCTAAGAAGGACCAGCCAGA |
|  | R- CCAGCAAACCTATCCACTCC |
| Caspase1 | F- AGATGCCAACCACTGAAAGG |
|  | R- GCATGATTCCCAACACAGGT |
